# Late Upper Palaeolithic hunter-gatherers in the Central Mediterranean: new archaeological and genetic data from the Late Epigravettian burial Oriente C (Favignana, Sicily)

**DOI:** 10.1101/692871

**Authors:** Giulio Catalano, Domenico Lo Vetro, Pier Francesco Fabbri, Swapan Mallick, David Reich, Nadin Rohland, Luca Sineo, Iain Mathieson, Fabio Martini

## Abstract

Grotta d’Oriente, a small coastal cave located on the island of Favignana (Sicily, Italy) is a key site for the study of the early human colonization of Sicily. The individual known as Oriente C was found in the lower portion of an anthropogenic deposit containing typical local Late Upper Palaeolithic (Late Epigravettian) stone assemblages. Two radiocarbon dates on charcoal from the deposit containing the burial are consistent with the archaeological context and refer Oriente C to a period spanning about 14,200-13,800 cal. BP. Anatomical features are close to those of Late Upper Palaeolithic populations of the Mediterranean and show strong affinity with Palaeolithic individuals of San Teodoro. Here we present new ancient DNA data from Oriente C. Our results, confirming previous genetic analysis, suggest a substantial genetic homogeneity among Late Epigravettian hunter-gatherer populations of Central Mediterranean, presumably as a consequence of continuous gene flow among different groups, or a range expansion following the Last Glacial Maximum (LGM).

## 1. Introduction

In the last few years, developments in sequencing techniques have enabled the generation of an unprecedented amount of genomic data from past populations. In particular, ancient genomes from Upper Palaeolithic and Mesolithic periods have made it possible to explore the early genetic makeup of hunter-gatherers of the (Jones et al., 2015; Fu et al., 2016; Hofmanová et al., 2016; Posth et al., 2016; Modi et al., 2017; Mathieson et al., 2018). Given its geographic location and the presence of Upper Palaeolithic and Mesolithic human remains belonging to at least 3 individuals (Oriente A, B and C: Mannino et al., 1972; Lo Vetro and Martini, 2006; Di Salvo et al., 2012; Mannino et al., 2012; Martini et al., 2012a), Grotta d’Oriente, on the island of Favignana (SW Sicily), is a key site for the study of the human colonization of Sicily during the Upper Palaeolithic (Lo Vetro and Martini, 2012). Paleogenetic and morphological studies on the Mesolithic individual Oriente B indicated a close proximity with Late Epigravettians of the Italian Peninsula (D’Amore et al., 2010; Mannino et al., 2012), while genome-wide single nucleotide polymorphism (SNP) data showed that the Upper Palaeolithic Oriente C clusters closely with other Western European Hunter-Gatherer (WHG) populations from Mesolithic and Late Palaeolithic Western Europe (Mathieson et al., 2018), confirming previous morphological analysis (Henke, 1989; Brewster et al., 2014). Here we generated new genome-wide data in order to refine the genetic affinities of Oriente C to other European hunter-gatherer populations. Our results and additional population genetic analyses provide insights into the origin and population structure of the hunter-gatherers that inhabited Europe during the Late Upper Palaeolithic and Mesolithic.

## 2. Archaeological setting

### 2.1. The site and its setting

Grotta d’Oriente is a small coastal cave located on the island of Favignana, the largest (∼20 km^2^) of a group of small islands forming the Egadi Archipelago, ∼5 km from the NW coast of Sicily (Fig. 1 A-B). The cave opens on the north-eastern slope of Montagna Grossa, at ∼40 m.a.s.l. (Fig. 1 C-D) The cave is formed of two distinct areas: a small chamber at the left of the entrance and a large gallery on the right (Martini et al., 2012b) (Fig. 1 E). Early excavations were conducted, without a strict methodology, in the small chamber in 1972 (Mannino, 1972; 2002). New excavations were performed in 2005 as a part of an interdisciplinary project carried out by the University of Florence and Museo e Istituto Fiorentino di Preistoria (Colonese et al., 2011, 2014, 2018; Craig et al., 2010; Lo Vetro and Martini, 2006; Martini et al., 2012a,b). A new trench was opened in 2005 next to the trench excavated in the 1970s and accurate recovery of materials and a microstratigraphic approach were followed. During the new excavations, a well-detailed archaeological sequence was documented (Fig. 1 F): the deposit, investigated up to a depth of about 2m, consists of 8 main sedimentological units (layers); five of which contain evidence of human frequentation of the cave during prehistory: Late Upper Palaeolithic (Layer 7), Early Mesolithic (Layer 6), Late Mesolithic or Early Neolithic (Layer 5) and Bronze Age (Layers 4-3). These cultural deposits were further divided into sublayers each corresponding to different paleosurfaces which are often characterized by hearths (more or less structured) and pits, and abundant artefacts and faunal remains (both terrestrial and marine). These sublayers (AMS radiocarbon data are in Table 1) are attributable to short-term episodes of human frequentation.

**Table 1.** Grotta d’Oriente, radiocarbon age for the stratigraphic sequence. 14C ages are reported as conventional and calibrated years BP IntCal13 (Reimer et al., 2013) in OxCal v4.3.

| Layer | Material | 14C yr BP | 14C yr cal BP<br>(68%) | 14C yr cal BP<br>(95%) | Cultural period | Reference |
| --- | --- | --- | --- | --- | --- | --- |
| 5A | Charcoal | 7040 ± 55 | 7940 – 7829 | 7969 – 7741 | Late Meso-Early Neolithic | Lo Vetro and Martini, 2006<br>Martini et al., 2012 |
| 6B | Charcoal | 8619 ± 65 | 9660 – 9530 | 9762 – 9485 | Early Mesolithic | Lo Vetro and Martini, 2006<br>Martini et al., 2012 |
| 6C | Charcoal | 8608 ± 65 | 9658 – 9526 | 9737 – 9480 | Early Mesolithic | Lo Vetro and Martini, 2006<br>Martini et al., 2012 |
| 6D | Charcoal | 8699 ± 60 | 9732 – 9551 | 9888 – 9542 | Early Mesolithic | Lo Vetro and Martini, 2006<br>Martini et al., 2012 |
| 7D | Charcoal | 12149 ± 65 | 14136 – 13932 | 14195 – 13791 | Upper Palaeolithic | Unpublished |
| 7E | Charcoal | 12132 ± 80 | 14107 – 13853 | 14198 – 13765 | Upper Palaeolithic | Lo Vetro and Martini, 2006<br>Martini et al., 2012 |

**Figure 1.**
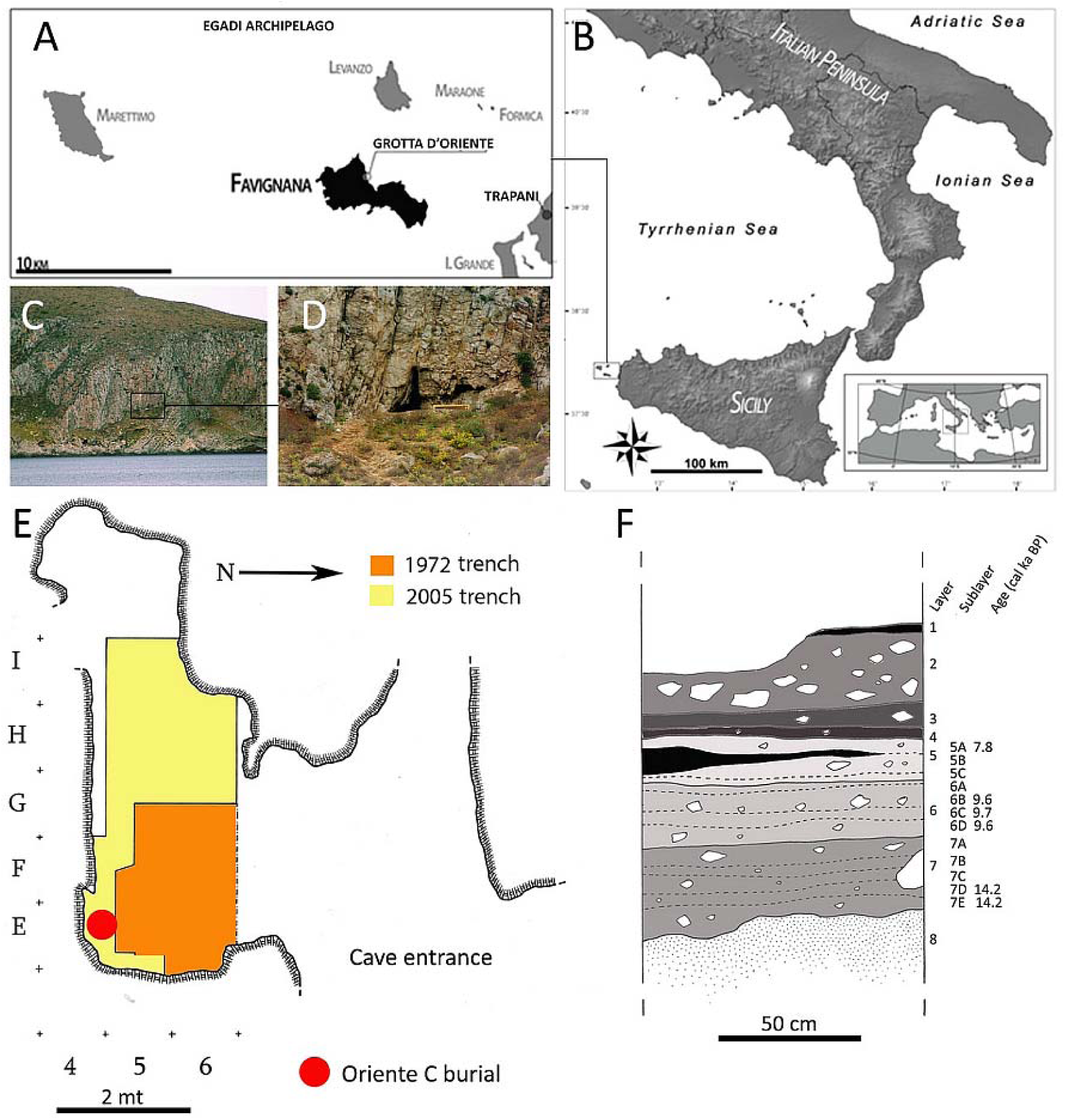
A-B) Geographic location of Grotta d’Oriente; C-D) the cave entrance on the slope of Montagna Grossa; E) excavation areas; F) stratigraphic sequence showing the layers and sublayers.

Below this sequence is a layer (Layer 8), containing Pleistocene fauna with no evidence of human activity. Above it are two historical levels (Layers 1-2), with scarce pottery remains, which have been largely reworked. In addition to the Palaeolithic burial Oriente C, which is the object of this study and was brought to light during the excavations in 2005 (Lo Vetro and Martini, 2006; Martini at al., 2012a), two other burials were discovered at Grotta d’Oriente in 1972: Oriente A (most likely a Late Palaeolithic adult male) and Oriente B (Mesolithic adult female) (Mannino, 2002; Mannino et al., 2012 and references therein).

### 2.2. The burial Oriente C

The Oriente C funereal pit opens in the lower portion of layer 7, specifically sublayer 7D. Two radiocarbon dates on charcoal from the sublayers 7D (12149±65 uncal. BP) and 7E, 12132±80 uncal. BP are consistent with the associated Late Epigravettian lithic assemblages (Lo Vetro and Martini, 2012; Martini et al., 2012b) and refer the burial to a period between about 14200-13800 cal. BP, when Favignana was connected to the main island (Agnesi et al., 1993; Antonioli et al., 2002; Mannino et al. 2014). Several attempts at direct radiocarbon dating on Oriente C were made by CEDAD laboratories (University of Lecce) but were unfortunately all unsuccessful.

The burial is located at the SW corner of the small chamber at the cave entrance, close to the rock wall. The skeleton is completely devoid of the lower limbs, large part of the pelvis, and the hands, because the burial was partly disturbed by two events: 1) during antiquity (in the Early Mesolithic), there was a small pit that originated from the top of the Layer 7 that partially affected the burial, perhaps removing the lower left skeletal elements; 2) in the early 1970’s, the trench excavated by G. Mannino intercepted the skeleton at the level of the pelvis as is clearly visible in Fig. 2.

**Figure 2.**
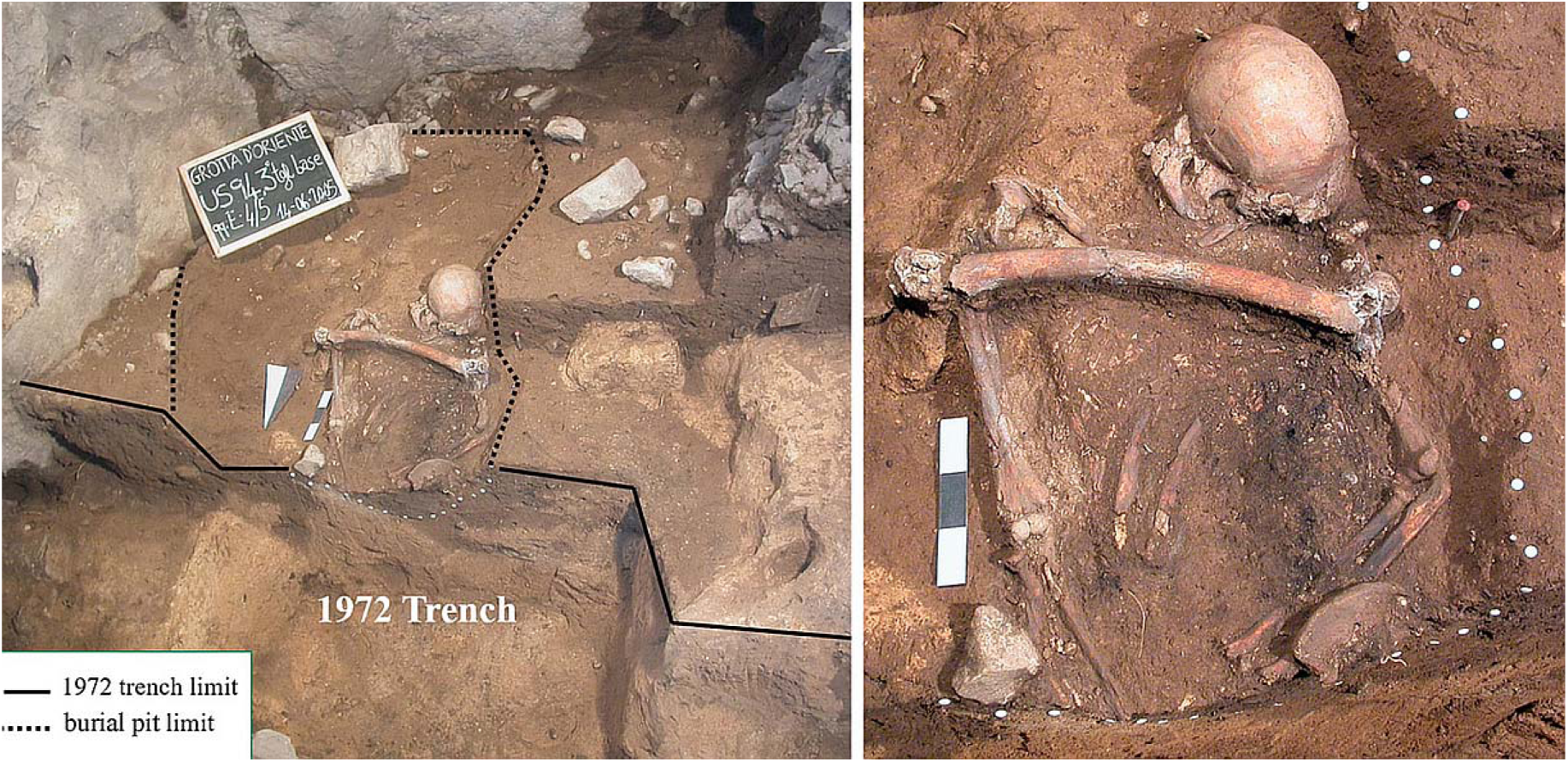
The burial Oriente C during the excavation: a wide view of the burial on the left, a close-up of the skeleton on the right.

In addition to these two events, a third one must be taken into account: a very peculiar feature of this burial is the presence of a femur (left) placed between the shoulders of Oriente C, on the thorax. The taphonomic data do not allow us to detect post-depositional disturbance of the skeleton that could have occurred in case of a reopening of the grave; the dislocations of some bones could therefore be attributed to post-burial movements inside the funerary pit. For this reason the femur could have been deposited during the interment and could have belonged to another individual (perhaps a relic).

However since the femur is compatible with the rest of the articulated skeleton (see below) we also need to consider the possibility that the femur belonged to Oriente C and resulted in a disturbance of the grave during the Palaeolithic which caused a dislocation of the lower bones after the decomposition of the corpse. This hypothesis could explain why G. Mannino did not notice the presence of Oriente C burial, which he would have to intercept with his excavation trench, because the skeleton was already devoid of the lower limbs (Mannino, 2002; Lo Vetro and Martini, 2006). If this event happened, it would have preceded the disturbance in the Mesolithic (the small pit opened at the top of layer 7A) as the grave was closed again after the deposition of the femur on the individual’s chest and sealed by the Palaeolithic deposit (layer 7C).

Beyond many disturbances and severe diagenetic phenomena, the preservation of the human remains at the site was very poor. To allow the recovery of human remains avoiding irreparable damage, during excavation it was necessary to consolidate them several times with abundant quantities of Paraloid B72, dissolved in acetone. Subsequently, the remains were removed and restored by gluing the parts after careful cleaning of the surfaces with acetone in order to remove the Paraloid film. Before any consolidation and restoration, during the burial excavation numerous bone fragments were recovered, and later determined by morphological examination to be most likely human.

Oriente C laid in dorsal decubitus oriented from South (the skull’s position) to North. The head rested on a large limestone chip with the face was turned slightly to the left. The right upper limb was extended on the side of the trunk, while the left one was flexed (about 120°) with its lower end placed on the lower abdomen. The bones of Oriente C curated in the Museo Fiorentino di Preistoria in Florence are the following: skull cap with fragmentary base; fragments of mandibular rami; lt. M^3^; dental fragments comprising a fragment of an upper molar larger than the M^3^. Rt. humerus: fragmentary lower third diaphysis; fragments of lo. epiphysis. Lt. Humerus: lower epiphysis. Rt. Radius: head. Lt. radius: up. epiphysis; fragments of diaphysis. Rt. ulna: upper 2/3 of diaphysis. Lt. ulna: up. and lo. epiphysis missing styloid process; fragmentary diaphysis. Lt. iliac bone: fragment comprising the ant. sup. Iliac spine. Lt. femur: missing lo. epiphysis. Lt. 3^rd^ metatarsal. All of these bones show the same colour (red-brownish) and degree of fossilization except the left ulna which is darker and the left radius which is nearly black because of a small fire lit in the grave (see above). The skull and mandible and left elbow were slightly displaced, upper right limb long bones were articulated, and the left iliac blade partially covered the flexed left forearm. These bones certainly belong to a single intentionally buried individual. A left femur was laid transversely above the upper part of the articulated skeleton, with the upper epiphysis on the left humerus (Fig. 2). No cut-marks or other traces linked to defleshing are evident on the femur. It is possible but not proven that the femur belongs to the same individual represented by the articulated bones (Lo Vetro and Martini, 2006). Several other human bones, often fragmentary, curated in the Museo Archeologico Regionale ‘‘Antonino Salinas’’ in Palermo (Mannino et al., 2012) were found in the Oriente cave during ’70 excavations: some of these bones could belong to Oriente C, especially the hand bones, but it is not possible at the moment to establish this.

The left humerus found in trench B does not belong to Oriente C as both humeruses are among the articulated bones curated in Florence; the distal left radius and right ulna from trench B are not represented in Florence and they could belong to Oriente C.

Age at death could not be accurately determined because of the lack of suitable anatomical parts. Nevertheless, we observe that rgw exocranial sutures are not fused and there is a beginning of fusion on the endocranial aspect of the obelic suture; the lt. M3 and the fragment of upper molar (an M1 or M2) are unworn, the six preserved long bone epiphysis (inferior right and left humerus; upper right and left radius; upper and lower left ulna) are completely fused to the diaphysis and do not show traces of osteoarthritis. We conclude that the individual probably was a young adult, maybe 25-30 years old. Oriente C lacks the diagnostic parts of the hip bones, but the long bone midshaft and epiphysis measurements – commonly used in sex determination of fragmentary human remains –indicated that the individual was most likely female (see Table S1 and Figure S1 in Supplementary material). This determination was later confirmed genetically (Mathieson et al., 2018). Stratigraphic and taphonomic features suggest that the funerary ritual of Oriente C consisted of a sequence of steps that can be summarized as follows:

*1-* *Excavation of the funerary pit*. The pit originates in the lower part of the layer 7 (sub-layer 7D) and affects the base of the layer 7 (sub-layer 7E) and the underlying layer 8 (sterile yellowish sands); It is shallow (about 25 cm) and has a flat bottom. The original mouth of the pit may have been obliterated in the case of a subsequent reopening of the pit (see step 5)The North portion of the pit was removed during the excavations in the 1970s.
*2-* *Deposition of the body*. The individual was placed into the grave, his skull resting almost on the western edge of the pit;
*3a-* *Burning action 1*. After the deposition, when the soft tissues of the body were probably still present, a low-heat fire was lit at the bottom of the grave, in direct contact with the body, in the area of the lower left hemithorax. The short and weak combustion left traces on the left forearm and deposited charcoal and ashy soil at the bottom of the pit. Following the hypothesis that the femur was placed on the shoulders of the deceased during the interment, this event must have occurred after the fire was already extinguished, since the femur lay on a thin layer of soil covering the charcoal and there are no traces of burning on the femur.
*3b-* *Burning action 2*. A second low-heat fire was lit to the right of the skull.
*4-* closing of the grave. The individual was definitively interred
*5-* *Possible reopening of the grave and deposition of a femur (if the femur belonged to Oriente C individual)*. A femur was placed between the shoulders of the body after the reopening of the grave. In this case the original mouth of the grave may had been partially obliterated and pit limits detected during the excavation may refer to the reopening of the burial. The reopening, if there was any, took place during the Late Upper Palaeolithic (top of layer 7D) since layer 7C, which covers the mouth of the pit, still refers to the Late Epigravettian.
*6-* *Deposition of stones*. Along the eastern edge of the grave, and also inside it, limestone blocks were deposited. Some of these blocks protruding from the pit were probably placed as a marker for the identification of the location of the burial.

The anatomical features of Oriente C are close to those of Late Upper Palaeolithic populations of the Mediterranean and show strong affinity with other Palaeolithic individuals of Sicily. As suggested by Henke (1989) and Fabbri (1995) the hunter-gatherer populations were morphologically rather uniform. This interpretation is further supported by the low or negligible *D*^*2*^ distance demonstrated by D’Amore et al. (2009) in the comparison between San Teodoro (individuals1-2-3-5-7) craniofacial morphometrics and other Upper Palaeolithic individuals. Like other Late Epigravettian burials in Sicily and Italy (Palma di Cesnola, 2006), Oriente C is a simple burial with little or no grave goods and personal ornaments. The only items in the pit were a pierced shell of *Cerithium* sp. (perhaps a clothing ornament) and a few small lumps of red ochre, next to the skull and the femoral head. Stable isotope analysis suggested a largely terrestrial diet with low-level consumption of marine foods which is comparable to other Late Upper Palaeolithic individuals from Sicily and Italy (Craig et al., 2010; Mannino et al.,2012).

## 3. Genetic analysis

A previous attempt at mitochondrial DNA analysis on a rib fragment of Oriente C, performed in 2006 by University of Rome ‘‘Tor Vergata” (Lo Vetro and Martini, 2006), was unsuccessful. The current ancient DNA analysis was done on a single long bone fragment that was not exposed to the substances used for consolidation and restoration (see above section 2.2). Sample preparation, DNA extraction and library construction were carried out in dedicated ancient DNA facilities in Boston as described in (Mathieson et al., 2018). To increase coverage compared to the previously reported data, we generated two additional libraries from the same extract, performed in-solution enrichment (“1240k”) and sequenced the product on an Illumina NextSeq500 using v.2 150 cycle kits for 2 × 76 cycles and 2 × 7 cycles. We merged these data with the data from the original library and made pseudo-haploid calls by selecting a single sequence at each single nucleotide polymorphism (SNP). The resulting dataset contained information on 288,223 SNPs covered at least once, compared to 61,547 in a previous publication (Mathieson et al.,2018), allowing for higher resolution analysis. To investigate the genetic affinities of Oriente C, we used the *qp3Pop* program from ADMIXTOOLS (Patterson et al., 2012) to compute *f*_*3*_*-*statistics and to estimate the amount of shared genetic drift between Oriente C and 98 published Mesolithic and Late Palaeolithic hunter-gatherers (with coverage at a minimum of 20,000 of 1240k positions) from 30 sites across Europe (Gamba et al.,2014; Gonzales-Fortes et al.,2017; Gunther et al.,2018; Haak et al.,2015;Jones et al.,2015; Lazaridis et al.,2014; Lipson et al.,2017; Mathieson et al.,2015,2018; Olalde et al.,2014). We used the *qpDstat* program to estimates *D*-statistics to test whether pairs of populations form a clade. The statistic *D(outgroup, A, B, C)* is zero if A is an outgroup to the clade (B,C), positive if A is closer to C, and negative if it is closer to B. For *D-* and *f*_*3*_-statistics, we estimated standard errors using the default block jackknife procedure implemented in ADMIXTOOLS (Patterson et al., 2012).

We confirmed the originally reported mitochondrial haplogroup assignment of U2’3’4’7’8’9. This haplogroup is present in both pre- and post-LGM populations, but is rare by the Mesolithic, when U5 dominates (Posth et al.2016). We further confirmed that the new genome-wide data was consistent with the original data by computing *D*-statistics (Patterson et al.,2012) of the form *D (Mbuti, X, Original Oriente data, Merged Oriente data)*. None of these statistics were significantly non-zero when X ranged over other European Mesolithic hunter-gatherers (maximum |Z| = 1.8 in 34 tests), and present-day French (Z = −0.35) and Sardinian (Z = −0.13) populations.

Lipson et al. (2018) (their supplementary Figure S5.1) and Villalba-Mouco et al. (2019) (their Figure 2A) showed that European Late Palaeolithic and Mesolithic hunter-gatherers fall along two main axes of genetic variation. Multidimensional scaling (MDS) of *f*_*3*_-statistics shows that these axes form a “V” shape (Fig. 3). At the point of the “V” lie the individuals that have been described as belonging to the “Western hunter-gatherer” (WHG) population, clustering closely with the 8,000 BP Loschbour individual (Lazaridis et al.2014). One arm represents a cline of ancestry that links WHG with “Eastern hunter-gatherer” (EHG) populations who carry ancestry related to the “Ancient North Eurasian” (ANE) population represented by the 24,000 BP Mal’ta individual (Raghavan et al., 2014). Along this cline lie Eastern European hunter-gatherer populations such as those from the Balkan Peninsula, present-day Ukraine (Mathieson et al., 2018), the Baltic (Jones et al., 2017; Mathieson et al., 2018) and Scandinavia (Haak et al., 2015, Gunther et al., 2018). The other arm of the “V” is a cline containing Late Upper Palaeolithic and Mesolithic individuals from Iberia (for example the individuals from El Mirón and La Braña), and Late Upper Palaeolithic individuals from Central Europe (Goyet and Hohle Fels). As shown by Fu et al. (2016), this cline reflects an ancestry contribution from a population related to the 35,000 BP Aurignacian Goyet Q116-1 individual. In this analysis, Oriente C falls at the tip of the “V”, at the extreme end of the WHG grouping.

**Figure 3.**
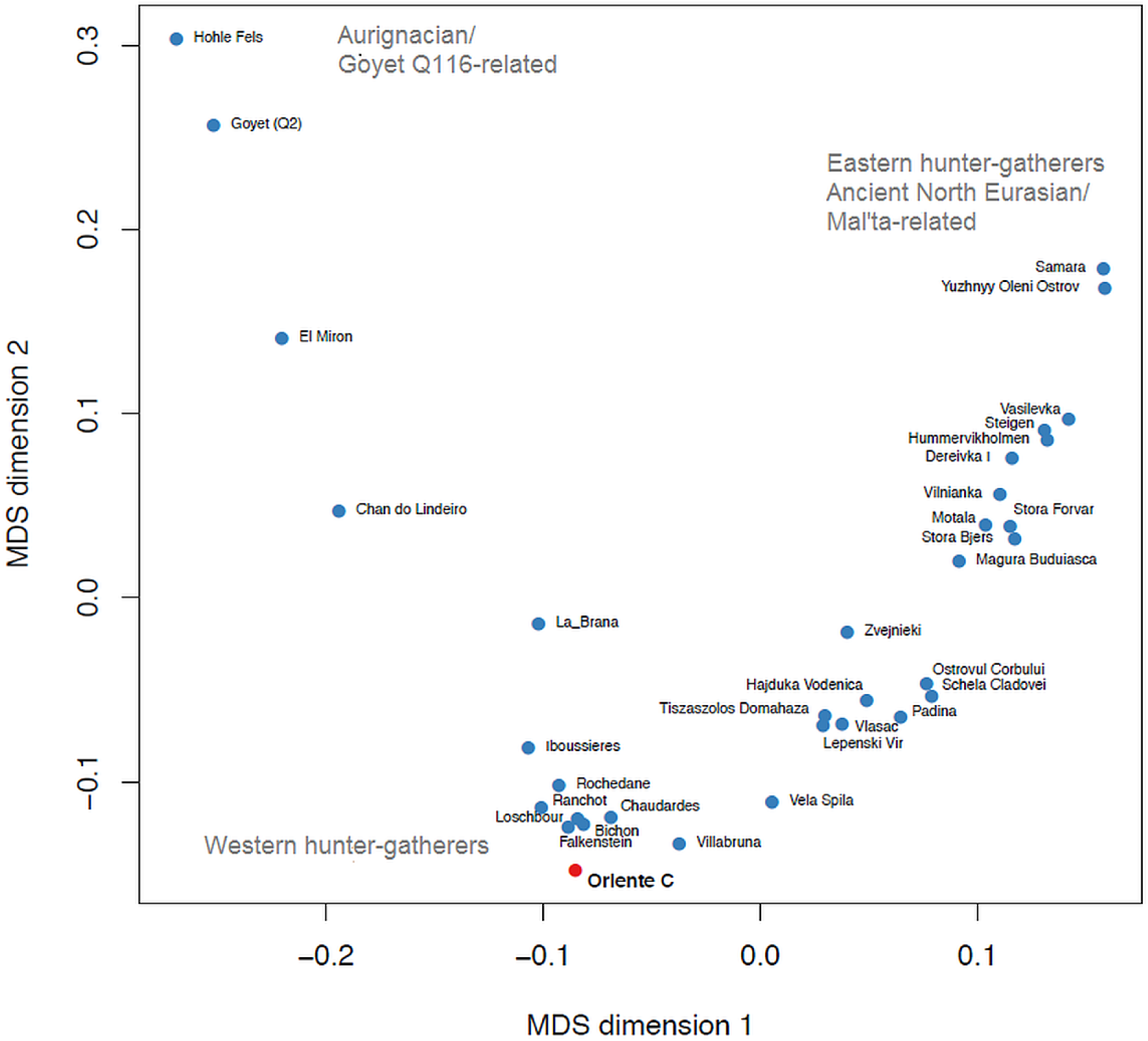
Multidimensional scaling of outgroup f_3_-statistics for Late Upper Palaeolithic and Mesolithic hunter-gatherers.

Focusing further on Oriente C, we find that it shares most drift with individuals from Northern Italy, Switzerland and Luxembourg, and less with individuals from Iberia, Scandinavia, and East and Southeast Europe (Fig. 4A-B). Shared drift decreases significantly with distance (Fig. 4C) and with time (Fig. 4D) although in a linear model of drift with distance and time as a covariate, only distance (p=1.3×10^-6^) and not time (p=0.11) is significant. Consistent with the overall E-W cline in hunter-gatherer ancestry, genetic distance to Oriente C increases more rapidly with longitude than latitude, although this may also be affected by geographic features. For example, Oriente C shares significantly more drift with the 8,000 year-old 1,400 km distant individual from Loschbour in Luxembourg (Lazaridis et al.,2014), than with the 9,000 year old individual from Vela Spila in Croatia (Mathieson et al.,2018) only 700 km away as shown by the D-statistic (Patterson et al.,2012) D (Mbuti, Oriente C, Vela Spila, Villabruna); Z=3.42. Oriente C’s heterozygosity was slightly lower than Villabruna (14% lower at 1240k transversion sites), but this difference is not significant (bootstrap P=0.12).

**Figure 4.**
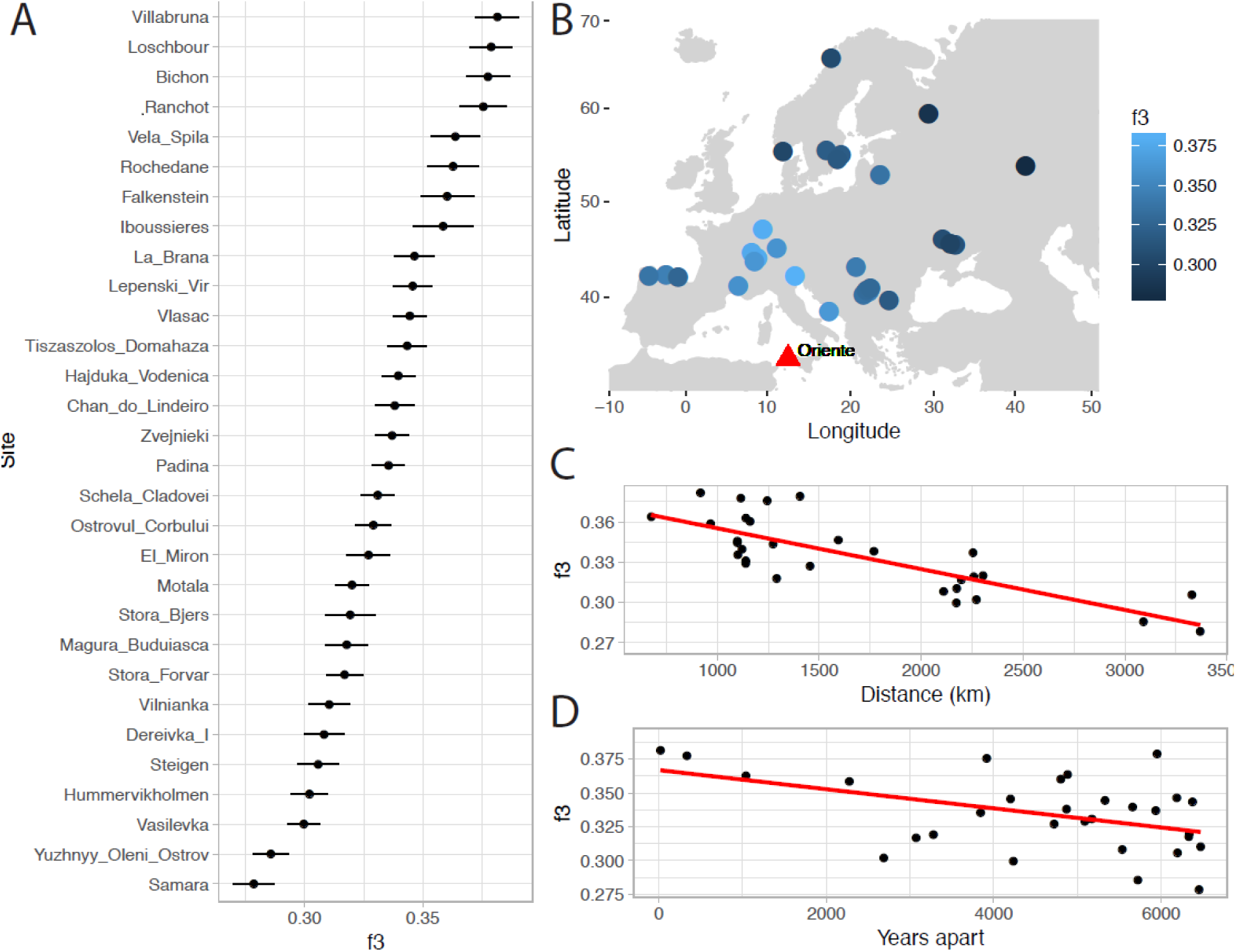
A) Shared drift, estimated using *f*_*3*_-statistics between Oriente C and 98 Mesolithic and Late Palaeolithic hunter-gatherers from 30 sites; B) The same statistic as in A plotted with geographic position; C) Decay of shared drift with distance from Oriente C; D) Decay of shared drift with absolute difference in date from Oriente C.

## 4. Discussion

Sicily falls within the area of expansion of the Epigravettian model widespread after the LGM in Mediterranean Europe, from Provence to the eastern Balkan border up to the Black Sea and the SW Anatolia (Fontana et al. in press). This “cultural province” is characterized by peculiar features which concern not only lithotechnics but also artistic production and burial customs. The Epigravettian (about 21.0-11.5 cal. ka BP) is a homogeneous cultural phenomenon despite the paleo-environmental differences occurring in a wide territory and some differentiations in resources exploitation strategies and human-environment interactions which were, perhaps, responsible for the appearance of regional variants. This homogeneous structure is also recognizable in Sicily despite the fact that the Late Epigravettian culture in the island presents a very specific local aspect, especially in lithic productions. The occurrence of cosmopolitan expressions and behaviours (iconographic languages, funeral rite) make Sicily the most continental of the Mediterranean islands with respect to material culture at the end of the Upper Palaeolithic.

The large number of archaeological sites related to the human frequentation of Sicily in the Late Upper Palaeolithic reveal that Late Epigravettian hunter-gatherer groups inhabited intensively the island during the Late Glacial period (Lo Vetro and Martini, 2012). The robust record of radiocarbon dates proves that they reached Sicily not before 15-14 ka cal. BP, several millennia after the LGM peak. In our opinion, in fact, the hypothesis about an early colonization of Sicily by Aurignacians (Laplace, 1964; Chilardi et al., 1996) must be rejected, on the basis of a recent re-interpretation of the techno-typological features of the lithic industries from Riparo di Fontana Nuova (Martini et al., 2007; Lo Vetro and Martini, 2012; on this topic see also Di Maida et al., 2019).

The Late Upper Palaeolithic burial Oriente C is a simple burial, and its sober ritual and the modality of deposition fit very well in the context of the Late Epigravettian burials of Sicily and Central-Southern Italy (Palma di Cesnola, 2006). Regarding the funerary ritual, an interesting issue concerns the occurrence of a femur placed on the shoulders of the individual and its possible belonging to the skeleton found in place.

Many sources of evidence indicate that the LGM may have had a major role in shaping the genetic and phenotypic variation of Upper Palaeolithic populations. A recent study based on complete mitochondrial genomes has revealed a genetic homogeneity between European hunter-gatherers. A significative predominance of the U lineage was detected with most of the sequences belonging to U5 haplotypes (Posth et al.2016). The finding of the haplogroup U2’3’4’7’8’9 in the Oriente C individual, previously recovered in the Upper Palaeolithic humans from Grotta Paglicci (Posth et al.,2016) provides additional evidence for the hypothesis that Epigravettian culture might have reached Sicily during the migration of Upper Palaeolithic groups from Southern Italy around the LGM (Palma di Cesnola, 2006; Lo Vetro and Martini, 2012; Mannino et al., 2012), which accords with the morphological similarity of Late Upper Palaeolithic and Early Mesolithic populations in the region (Henke, 1989; Brewster et al., 2014). The find of genetic similarity of Oriente C with Late Upper Palaeolithic and Mesolithic individuals from Northern Italy (i.e. Villabruna) and Central Europe (i.e. Bichon, Loschbour) (Fig. 3) is also in line with previous studies according to which Sicilian hunter-gatherers were found to be morphologically closely related to Late Epigravettians of the Italian Peninsula and continental Europe (Fabbri, 1995; D’Amore et al., 2010).

## 5. Conclusion

These analyses have implications for understanding the origin and diffusion of the hunter-gatherers that inhabited Europe during the Late Upper Palaeolithic and Mesolithic. Our findings indicate that Oriente C shows a strong genetic relationship with Western European Late Upper Palaeolithic and Mesolithic hunter-gatherers, suggesting that the “Western hunter-gatherers” was a homogeneous population widely distributed in the Central Mediterranean, presumably as a consequence of continuous gene flow among different groups, or a range expansion following the LGM.

The DNA study of Oriente C is particularly relevant to studying the peopling of the Central Mediterranean by Anatomically Modern Humans after the LGM. The data support the hypothesis that hunter-gathering groups arrived in Sicily from the Italian peninsula, confirming results derived from anatomical studies on human fossil remains of Grotta di San Teodoro and from the stone assemblages whose features fit in the panorama of the Late Epigravettian of Southern Italy.

## Data accessibility

The 1240k capture sequencing data for Oriente C (merged new and existing data) has been deposited in the European Nucleotide Archive (https://www.ebi.ac.uk/ena) under accession number PRJEB33231.

## Acknowledgments

The authors thank and remember with affection their friend and colleague Sebastiano Tusa, prematurely died in the plane crash of the Ethiopian Airlines in 2019. Dr. Tusa did so much for understanding the prehistoric heritage of Sicily.

## Authors’ Contributions

GC and DLV designed the paper, DLV and FM conducted the excavation at Grotta d’Oriente, PFF conducted the anthropological study of the human remains, DLV, FM, PFF provide information about the taphonomy of the burial, LS collected the sample, IM, SM, NR and DR conducted the DNA analysis, GC, IM, DLV, PFF wrote the manuscript with input from all co-authors.

## References

Agnesi, V., Macaluso, T., Orrù, P., Ulzega A., 1993. Paleogeografia dell’Arcipelago delle Egadi (Sicilia) nel Pleistocene superiore-Olocene, Naturalista siciliano, S. IV, XVII (1-2), pp. 3–22.

Antonioli, F., Cremona, G., Immordino, F., Puglisi, C., Romagnoli, C., Silenzi, S., Valpreda, E., Verrubbi, V., 2002. Newdata on the Holocenic sea-level rise in NW Sicily (Central Mediterranean Sea), Global and Planetary Change, pp. 121–140.

Brewster, C., Meiklejohn, C., von Cramon-Taubadel, N., Pinhasi, R., 2014. Craniometric analysis of European Upper Palaeolithic and Mesolithic samples supports discontinuity at the Last Glacial Maximum. Nature Communications 5:4094. https://doi.org/10.1038/ncomms5094.

Chilardi, S., Frayer, D.W., Gioia, P., Macchiarelli, R., Mussi, M., 1996. Fontana Nuova di Ragusa (Sicily, Italy): southernmost Aurignacian site in Europe. Antiquity 70: 553–563.

Colonese, A.C., Zanchetta, G., Drysdale, R.N., Fallick, A.E., Manganelli, G., Lo Vetro, D.,Martini, F., Di Giuseppe, Z., 2011. Stable isotope composition of latePleistocene-Holocene Eobania vermiculata (Müller, 1774) (Pulmonata, Stylommatophora)shells from the Central Mediterranean basin: data from Grottad’Oriente (Favignana, Sicily). Quaternary International 244, 76–87.https://doi.org/10.1016/j.quaint.2011.04.035.

Colonese, A.C., Lo Vetro, D., Martini, F., 2014. Holocene coastal change and intertidal mollusc exploitation in the central Mediterranean: variations in shell size and, morphology at Grotta d’Oriente (Sicily). Archaeofauna 23, 181–192.

Colonese, A.C., Lo Vetro, D., Landini, W., Di Giuseppe, Z.,Hausmann, N., Demarchi, B., d’Angelo, C., Leng, M. J., Incarbona, A., Whitwood, A.C., Martini, F., 2018. Late Pleistocene-Holocene coastal adaptation in central Mediterranean: Snapshots from Grotta d’Oriente (NW Sicily), Quaternary International, 493, pp. 114–126. http://dx.doi.org/10.1016/j.quaint.2018.06.018.

Craig, O.E., Biazzo, M., Colonese, A.C., Di Giuseppe, Z., Martinez-Labarga, C., Lo Vetro, D., Lelli, R., Martini, F., Rickards, O., 2010. Stable isotope analysis of Late Upper Palaeolithic humans and fauna remains from Grotta del Romito (Cosenza), Italy. Journal of Archaeological Science 37, 2504–2512.

D’Amore, G., Di Marco, S., Tartarelli, G., Bigazzi, R., Sineo, L., 2009. Late Pleistocene human evolution in Sicily: Comparative morphometric analysis of Grotta di San Teodoro craniofacial remains. J Hum Evol 56:537–550.

D’Amore, G., Di Marco, S., Di Salvo, R., Messina, A., Sineo., L, 2010. The early peopling of Sicily: evidence from the Mesolithic skeletal remains from Grotta d’Oriente. Annals of Human Biology, 37, pp. 403–426.

Di Salvo, R., Mannino, G., Mannino, M.A., Schimmenti, V., Sineo, L., Thomas, K.D., 2012. Le sepolture della Grotta d’Oriente (Favignana). Atti della XLI Riunione Scientifica dell’Istituto Italiano di Preistoria e Protostoria: Dai Ciclopi agli Ecisti: società e territorio nella Sicilia preistorica e protostorica: 341–351.

Fabbri, P.F., 1995. Dental anthropology of the Upper Palaeolithic sample from San Teodoro and inferences on the peopling of Italy. Zeitschrift für Morphologie und Anthropologie, 80 311–327.

Fu, Q. et al., 2016. The genetic history of Ice Age Europe. Nature. 534(7606): 200–205. https://doi.org/10.1038/nature17993.

Gamba, C. et al., 2014. Genome flux and stasis in a five millennium transect of European prehistory. Nature Communications 5. https://doi.org/10.1038/ncomms6257.

Gonzalez-Fortes, G. et al., 2017. Paleogenomic Evidence for Multi-generational Mixing between NeolithicFarmers and Mesolithic Hunter-Gatherers in the Lower Danube Basin. Current biology: CB 27, 1801–1810. https://doi.org/10.1016/j.cub.2017.05.023.

Gunther, T. et al., 2018. Population genomics of Mesolithic Scandinavia: Investigating early post glacial migration routes and high-latitude adaptation. PloS Biol 16, e2003703. https://doi.org/10.1371/journal.pbio.2003703.

Haak, W. et al., 2015. Massive migration from the steppe was a source for Indo-European languages in Europe. Nature 522, 207–211. https://doi.org/10.1038/nature14317.

Henke, W., 1989. Biological distances in Late Pleistocene and Early holocene human population in Europe. In “People and Culture in change”.I. Hershkovitz (Ed.). Proc. of the 2nd Symp. on Up. Paleol., Mesol. and Neol. populat. of Europe and the Medit. Bas., Tel Aviv, Sept. 6-10, 1987. BAR Int. Ser. 508(ii):541–563.

Hofmanová, Z. et al., 2016. Early farmers from across Europe directly descended from Neolithic Aegeans. PNAS June 21, 2016 113 (25) 6886–6891. https://doi.org/10.1073/pnas.1523951113.

Jones, E.R. et al., 2015. Upper Palaeolithic genomes reveal deep roots of modern Eurasians. Nature Communications 6. https://doi.org/10.1038/ncomms9912.

Jones, E.R., et al., 2017. The Neolithic Transition in the Baltic Was Not Driven by Admixture with Early European Farmers. Curr Biol. 2185–2193. https://doi.org/10.1016/j.cub.2016.12.060.

Laplace G. 1964, Les subdivisions du leptolithique italien. Étude de typologie analytique. Bullettino di Paletnologia Italiana. 1964; 73: 25–63.

Lazaridis, I. et al., 2014. Ancient human genomes suggest three ancestral populations for present-day Europeans. Nature 513, 409–413. https://doi.org/10.1038/nature13673.

Lipson, M. et al., 2017. Parallel palaeogenomic transects reveal complex genetic history of early European farmers. Nature 551, 368–372. https://doi.org/10.1038/nature24476.

Lo Vetro, D., Martini, F., 2006. La nuova sepoltura epigravettiana di Grotta d’Oriente (Favignana, Trapani). In:Martini, F. (Ed.): 3. La cultura del Morire nelle società preistoriche e protostoriche italiane. Studio interdisciplinare dei dati e loro trattamento informatico. Origines, Progetti, vol. 1. Istituto Italiano di Preistoria e Protostoria, Firenze: 58–66.

Lo Vetro, D., Martini, F., 2012. Il Paleolitico e il Mesolitico in Sicilia. In: Atti XLI Riunione Scientifica IIPP, “Dai Ciclopi agli Ecisti: società e territorio nella Sicilia preistorica e protostorica”. San Cipirello, Italy, pp. 19–48.

Mannino, G., 1972. Grotta d’Oriente. Rivista di Scienze Preistoriche XXVII (2): 470.

Mannino, G., 2002. La Grotta d’Oriente di Favignana (Egadi, Sicilia). Risultati di un sondaggio esplorativo. Quaderni del Museo Archeologico Regionale Antonio Salinas 8: 9–22.

Mannino, M.A., Catalano, G., Talamo, S., Mannino, G., Di Salvo, R., Schimmenti, V.,Lalueza-Fox, C., Messina, A., Petruso, D., Caramelli, D., Richards, M.P., Sineo, L., 2012. Origin and diet of the prehistoric hunter-gatherers on the Mediterranean Island of Favignana (Egadi Islands, Sicily). PLoS One 7 (11), e49802. http://dx.doi.org/10.1371/journal.pone.0049802.

Mannino, M.A., Thomas, K.D., Crema, E.R., Leng, M.J., 2014. A matter of taste? Mode and periodicity of marine mollusc exploitation on the Mediterranean island of Favignana (Egadi Islands, Italy) during its isolation in the early Holocene. Archaeofauna: International Journal of Archaeozoology 23, 133–147.

Martini, F., Lo Vetro, D., Colonese, A.C., De Curtis, O., Di Giuseppe, Z., Locatelli, E., Sala, B. 2007, L’Epigravettiano Finale in Sicilia. In: Martini, F, editor. L’Italia tra 15.000 e 10.000 anni fa. Cosmopolitismo e regionalità nel Tardoglaciale. Firenze: Museo e Istituto Fiorentino di Preistoria “Paolo Graziosi”: 209–254.

Martini, F., Lo Vetro, D., Borrini, M., Bruno, S., Mallegni, F., 2012a. Una nuova sepoltura dalla Grotta di Oriente (Favignana, Trapani). Scavi 2005. Atti della XLI Riunione Scientifica dell’Istituto Italiano di Preistoria e Protostoria: Dai Ciclopi agli Ecisti: società e territorio nella Sicilia preistorica e protostorica: 333–340.

Martini F., Lo Vetro D., Colonese A.C., Cilli C., De Curtis O., Di Giuseppe Z., Giglio R., Locatelli E., Sala B., Tusa S., 2012b. Primi risultati sulle nuove ricerche stratigrafiche a Grotta d’Oriente (Favignana, Trapani). Scavi 2005. Atti della XLI Riunione Scientifica dell’Istituto Italiano di Preistoria e Protostoria: Dai Ciclopi agli Ecisti: società e territorio nella Sicilia preistorica e protostorica: 319–332.

Mathieson, I. et al., 2015. Genome-wide patterns of selection in 230 ancient Eurasians. Nature 528, 499–503.https://doi.org/10.1038/nature16152.

Mathieson, I. et al., 2018. The genomic history of southeastern Europe. Nature 555, 197–203. https://doi.org/10.1038/nature25778.

Modi, A. et al., 2017. Complete mitochondrial sequences from Mesolithic Sardinia. Sci Rep; 7: 42869. https://doi.org/10.1038/srep42869.

Olalde, I. et al., 2014. Derived immune and ancestral pigmentation alleles in a 7,000-year-old Mesolithic European. Nature 507, 225–228.https://doi.org/10.1038/nature12960.

Palma di Cesnola, A., 2006. L’Aurignacien et le Gravettien ancient de la grotte Paglicci au Mont Gargano. L’Anthropologie 110: 355–370.

Patterson, N., Moorjani, P., Luo, Y., Mallick, S.,Rohland, N.,Zhan, Y., Genschoreck, T.,Webster, T., Reich, D., 2012. Ancient admixture in human history. Genetics 192, 10651093. https://doi.org/10.1534/genetics.112.145037.

Posth, C. et al., 2016. Pleistocene Mitochondrial Genomes Suggest a Single Major Dispersal of Non-Africans and a Late Glacial Population Turnover in Europe. Current Biology 26, 827–833. https://doi.org/10.1016/j.cub.2016.01.037.

Raghavan, M, et al., 2014. Upper Palaeolithic Siberian genome reveals dual ancestry of Native Americans. Nature 505(7481):87–91. https://doi.org/10.1038/nature12736.

Reimer, P.J. et al., 2013. IntCal13 and Marine13 radiocarbon age calibration curves 0–50,000 Years cal BP. Radiocarbon 55, 1869–1887.

Villalba-Mouco, V. et al., 2019. Survival of Late Pleistocene Hunter-Gatherer Ancestry in the Iberian Peninsula. Current Biology 29, 1169–1177. https://doi.org/10.1016/j.cub.2019.02.006

